# ‘Candidatus Phytoplasma platensis’, a novel taxon associated with daisy (Bellis perennis) virescence and related diseases in South America

**DOI:** 10.1101/564807

**Authors:** Franco Daniel Fernández, Ernestina Galdeano, Luis Rogelio Conci

## Abstract

Bellis perennis virescence (BellVir) phytoplasma affects ornamental daisies in Argentina. It has been previously classified within the X-disease group, subgroup III-J, which is one of the most important and widely distributed in South America, affecting diverse plant hosts. In this study, we compared 16S rRNA, ribosomal proteins rpIV and rps3, secA and immunodominant proteins imp and idpA genes of BellVir phytoplasma with previously described ‘*Candidatus* Phytoplasma’ species. The 16S rRNA gene of strain BellVir shared less than 97.5% with all previously described ‘*Ca*. Phytoplasma’ taxa except for ‘*Ca*. Phytoplasma pruni’. According to the recommended rules for the description of novel taxa within ‘*Ca*. Phytoplasma’, it should be considered as ‘*Ca*. P. pruni’-related strain. However, multilocus analysis showed further molecular diversity that distinguished BellVir phytoplasma from ‘*Ca*. Phytoplasma pruni’. Besides, BellVir phytoplasma and 16SrIII-J related strains have a geographical distribution restricted to South America, where ‘*Ca*. P.pruni’ has not been detected. Two insect vectors have been reported to transmit 16SrIII-J phytoplasmas, which have not been found to transmit ‘*Ca*. Phytoplasma pruni’. Having a wide host range, they have not been detected in *Prunus persica*. Therefore, based on multilocus sequence analyses, specific vector transmission and geographical distribution, we propose the recognition of the novel phytoplasma species ‘*Ca*. Phytoplasma platensis’, within the X-disease clade, with Bellis perennis virescence phytoplasma as the reference strain.

---

The provisional genus ‘*Candidatus* Phytoplasma’, formally described in 2004, includes plant pathogenic non-helical *Mollicutes* that inhabit plant phloem and insects [1]. Phytoplasmas have been reported worldwide causing diseases in more than 1000 plant species [2]. In plant hosts they colonize mainly sieve tube elements but can reach parenchymatic companion cells. They are transmitted among plant hosts by sap-sucking insects, such as leafhoppers and psyllids, in a persistent manner and could be detected in the insects’ gut, hemolymph, salivary glands and other organs [3]. Symptomatic plants can show stunting, die back, abnormal shoot proliferation (witches’ broom), leaf yellowing or reddening, reduced leaf size and deformation. Other symptoms include flower sterility, phyllody and virescence [4, [5].

The provisional status of *Candidatus* responds to the difficulty of obtaining pure cultures, and consequently phenotypic description. Instead, conserved gene sequence analysis has been used for the detection and classification of phytoplasma strains. Based on 16S rRNA gene sequence and RFLP analysis phytoplasmas have been classified into 16Sr groups and subgroups [6, [7, [8]. So far, more than thirty groups have been described, most of which host at least one ‘*Ca*. Phytoplasma species’ [9, [10]. Further distinction can be achieved by including biological features such as plant hosts, symptoms, insect vectors, geographical distribution, and multilocus sequence analysis [11–14]. According to the recommended rules for the description of novel taxa within ‘*Ca*. Phytoplasma’, a new species should refer to a unique 16S rRNA gene sequence with ≤97.5% similarity with a previously described ‘*Ca*. Phytoplasma species’. However, a strain that has higher sequence similarity could be described as a new species if it has clearly different biological characteristics including host range and vector transmission, and significative molecular diversity [1].

*Bellis perennis* virescence phytoplasma (BellVir) (Figure S1) was first reported in 2013 affecting ornamental daisies in Argentina. Based on 16S rRNA and ribosomal protein genes analyses it has been classified within group 16SrIII or X-disease group, subgroup 16SrIII-J [15]. Chayote witches’ broom (ChWBIII) phytoplasma was the first 16SrIII-J strain detected in Brazil [16]. After that, subgroup 16SrIII-J phytoplasmas have been reported in various cultivated, domesticated and wild plant hosts in South America, mainly Argentina, Southern Brazil and Chile [15, 17–21]. Infected plant hosts showed diverse symptoms including leaf size and general growth reduction, internode shortening and shoot proliferation, color changing, phyllody and virescence. The X-disease or 16SrIII is one of the most numerous and diverse groups, composed by more than 25 subgroups. Up to now, one ‘*Ca*. Phytoplasma’ species has been described (‘*Ca*. Phytoplasma pruni’) that includes strains of group 16SrIII, with Peach X-disease phytoplasma from subgroup 16SrIII-A as reference strain (PX11CT1, JQ044392). Phytoplasmas from other 16SrIII subgroups have been considered as related strains since they show differences in the oligonucleotide unique regions of the rRNA gene [22], However, the high diversity within 16SrIII group led the authors to suggest that the analyzed 16SrIII group phytoplasma lineages might represent at least two, or even three different phytoplasma taxa..

*Bellis perennis* phytoplasma (BellVir, 16SrIII-J) [15] was selected as reference strain for molecular analyses. The phytoplasma was first transmitted from infected daisies to healthy periwinkle (*Catharanthus roseus* (L.) G. Don.) using *Cuscuta subinclusa*. Once the infection was established in periwinkle, the strain was perpetuated by periodical grafting. Typical symptoms of phyllody, virescence, leaf size reduction and yellowing were observed in infected plants 3 to 6 months after grafting (Figure S1). For PCR amplifications, genomic DNA was extracted from BellVir infected periwinkle leaves and petioles by the CTAB technique [23]. Phytoplasma 16S rRNA gene was amplified by PCR using primers P1/P7 [24] according to previously described protocol [15]. Partial secA gene amplification (∼0.80kb) was performed using SecAfor1/SecArev3 primer pair following the conditions proposed by [25]. For immunodominant proteins, we designed specific primers based on the genomic information available in public database. So far, only one draft genome has been described for subgroup 16SrIII-J [26] which belongs to Vc33 phytoplasma isolate from periwinkle. Annotation pipeline led us to identify imp and idpA gene sequences [27]. Two new sets of primers impXd-Fw1/impXd-Rv1 and idpAXd-Fw1/idpA-Rv1 were designed in order to amplify by PCR a genomic fragment containing the complete sequence of imp and idpA genes, respectively (Table S3, Supplementary material). The amplicons were purified and cloned as described previously [28]. Three clones for each isolate were bidirectionally sequenced using an automated DNA Sanger sequencer (Unidad Genómica, Instituto de Biotecnología-Instituto Nacional de Tecnología Agropecuaria, Argentina). Final consensus sequences (3X coverage) were assembled using the Geneious R10 software and deposited in the GenBank nucleotide database. The phylogenetic reconstruction for each gene was performed using Maximum Likelihood method from the MEGA 6 software package [29].

## BellVir represents a novel taxon for the provisional genus ‘Candidatus Phytoplasma’

The signature sequence (5’-CAAGAYBATKATGTKTAGCYGGDCT-3’) characteristic of the provisional genus ‘*Candidatus* Phytoplasma’ is contained in BellVir’s 16S rRNA gene (5’-CAAGACTATGATGTGTAGCTGGACT-3’) (263-287). The 16S rRNA gene (MK135798) of strain BellVir shared less than 97.5% with corresponding fragments of the 16S rRNA genes from all previously described ‘*Ca*. Phytoplasma’ taxa except for ‘*Ca*. Phytoplasma pruni’, with 98.79-98.87% nucleotide sequence identity (Table 1). However, multilocus analysis showed further molecular diversity that distinguished BellVir phytoplasma from ‘*Ca*. Phytoplasma pruni’. Besides, BellVir phytoplasma and 16SrIII-J related strains have a geographical distribution restricted to southern South America and have particular biological characteristics. Having a wide host range that includes *Bellis perennis* (used as reference strain), *Allium sativum, Cucurbita maxima, Coffea arabica, Solanum lycopersicum, S. melongea, Helianthus annus, Sechium edule, Brassica oleracea, Beta vulgaris, Fragaria* x *annanasa, Lactuca sativa, Manihot sculenta* and *Prunus avium* (cherry) among others[16, [20, 29–31, [33], phytoplasmas from this subgroup have not been detected in *Prunus persica* (peach). As regards vector transmission, two insect vectors have been reported to transmit 16SrIII-J phytoplasmas, *Paratanus exitiosus* and *Bergallia valdiviana*, which have not been found to transmit ‘*Ca*. Phytoplasma pruni’ [19, [34]. ‘*Ca*. Phytoplasma pruni’ has not been detected until now in South American countries and, if 16SrIII-J phytoplasmas were considered related to it, attempts to regulate the pathogen introduction into these countries would be very difficult to accomplish.

The phylogenetic tree based on 16S rDNA sequences of BellVir phytoplasma and known ‘*Ca*. phytoplasma species’ showed that BellVir clustered with ‘*Ca*. Phytoplasma pruni’ but separated into an independent branch within the cluster (Figure 1). A broader phylogenetic analysis showed that ‘*Ca*. Phytoplasma platensis’ and ‘*Ca*. Phytoplasma pruni’ strains conform two well defined clades (Figure S2, supplementary material). Previous works had shown the same topology, separating 16SrIII-J phytoplasmas from other 16SrIII subgroups [15, [18, [21]. When the unique signature regions that distinguish ‘*Ca*. Phytoplasma pruni’ where examined in BellVir’s 16S rRNA gene sequence, 7 out of the 13 unique regions showed at least one nucleotide difference between them (Figure S3, supplementary material).

**Figure 1:**
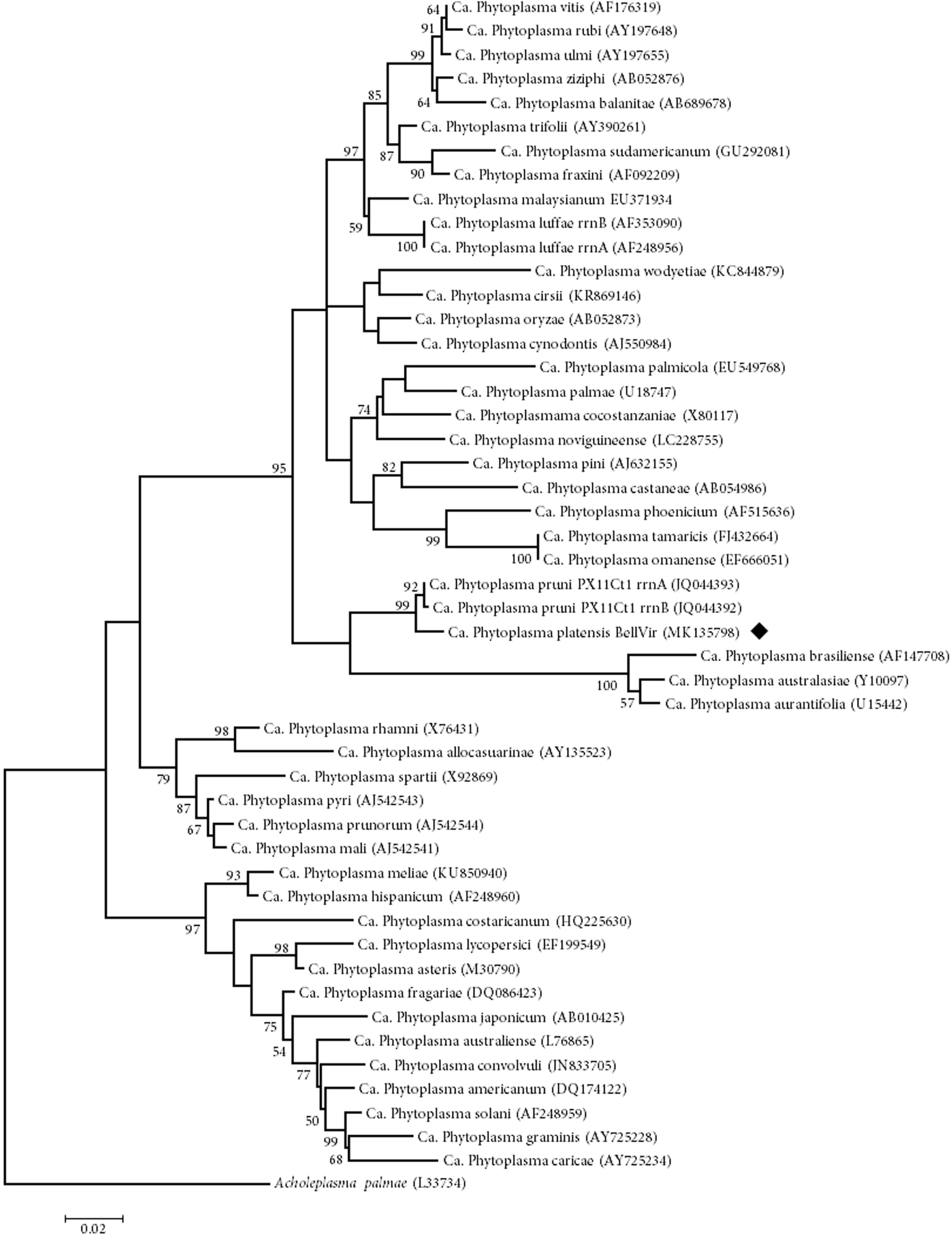
Phylogenetic tree inferred from 16S rRNA gene sequence analysis using the Maximum Likelihood method implemented in the Molecular Evolutionary Genetics Analysis program (MEGA 6). *Acholeplasma palmae* was used as outgroup. The numbers on the branches are bootstrap (confidence) values (expressed as percentage of 1000 replicates). GenBank accession number for each taxon is given between parentheses. The final tree includes reference strain of 48 previously described or incidentally cited as ‘Ca. Phytoplasma species’ and ‘Ca. Phytoplasma platensis’ (marked with black diamond). Bar, number of nucleotide substitutions per site.

### Multilocus sequence analyses differentiate ‘*Candidatus* Phytyoplasma platensis’ from ‘*Candidatus* Phytoplasma pruni’

Other genomic regions were examined to determine molecular diversity of BellVir comparing with previously described ‘*Ca*. Phytoplasma pruni’. Sequences of ribosomal protein genes rpIV and rps3 of BellVir showed >98.5% identity with related strains and 96.5% with ‘*Ca*. Phytoplasma pruni’. RFLP patterns generated by restriction enzymes *Alu*I and *Dra*I distinguished BellVir from ‘*Ca*. Phytoplasma pruni’ [15]. The phylogenetic tree showed that BellVir and related phytoplasmas integrate a cluster separated from ‘*Ca*. Phytoplasma pruni’ and related strains (Figure 2). Multiple alignment of ribosomal proteins gene sequences revealed the presence of 17 SNPs that distinguish ‘*Ca*. Phytoplasma platensis’ strains from the ‘Ca. Phytoplasma pruni’ strains (Table S2, supplementary material). Similar results were obtained when secA gene was analyzed, and the amino acid sequence corresponding to BellVir secA protein (MK140657) showed 99.4% identity with Vac33 strain (LLKK01000003) while both of them shared a maximum identity of 96.6% with ‘*Ca*. Phytoplasma pruni’ strains. The resulting tree had the same topology as the generated by 16S rRNA and ribosomal protein genes, supporting the separation of BellVir phytoplasma (Figure 3). Immunodominant proteins would not be correlated with that of 16Sr DNA; however, imp is a gene well conserved over a wide range of phytoplasmas and can reflect in between phytoplasma differences regarding host range and vector transmission [35]. Phytoplasmas immunodominant proteins have been classified into three distinct types: (i) immunodominant membrane protein (Imp); (ii) immunodominant membrane protein A (IdpA); and (iii) antigenic membrane protein (Amp) [36]. BellVir phytoplasma has the same type of immunodominant membrane proteins as all 16SrIII-group phytoplasmas since both imp and idpA genes could be amplified [37]. BellVir imp amino acid sequence had 97.7% identity with 16SrIII-J phytoplasma Vac33, and 58.3-61% identity with ‘*Ca*. Phytoplasma pruni’ related strains, which showed 97.7-79.6% identity among them. The phylogenetic tree constructed with imp aa sequences of 16SrIII phytoplasmas clearly separated BellVir from ‘*Ca*. Phytoplasma pruni’ and related strains (Figure S4, supplementary material). Similar situation arise within the idpA protein, since the highest identity occurs with Vac33 phytoplasma (87.4%) and 66.26-69.51% identity *Ca*. Phytoplasma pruni’ related strains, which showed 99.65-63.61% identity among them. The topology of the phylogenetic tree constructed with idpA aa sequences resembled those of imp, and showed once again a clear separation of ‘*Ca*. Phytoplasma platensis’ and ‘*Ca*. Phytoplasma pruni’ clades.

**Figure 2:**
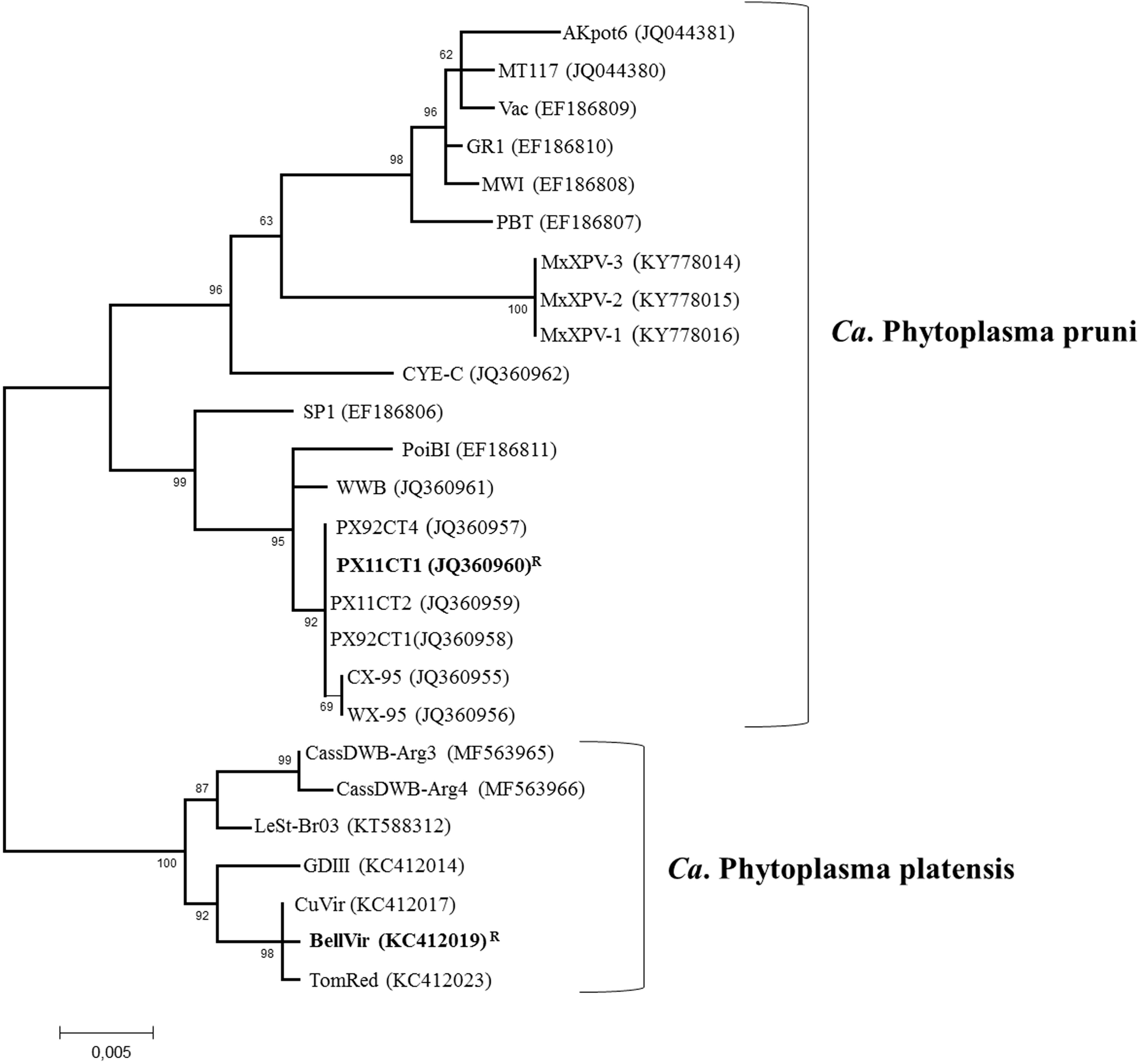
Phylogenetic tree inferred from rpIV and rps3 genes sequence analysis using the Maximum Likelihood method implemented in the Molecular Evolutionary Genetics Analysis program (MEGA 6). The numbers on the branches are bootstrap (confidence) values (expressed as percentage of 1000 replicates). GenBank accession number for each taxon is given between parentheses. R: reference strains. Bar, number of nucleotide substitutions per site.

**Figure 3:**
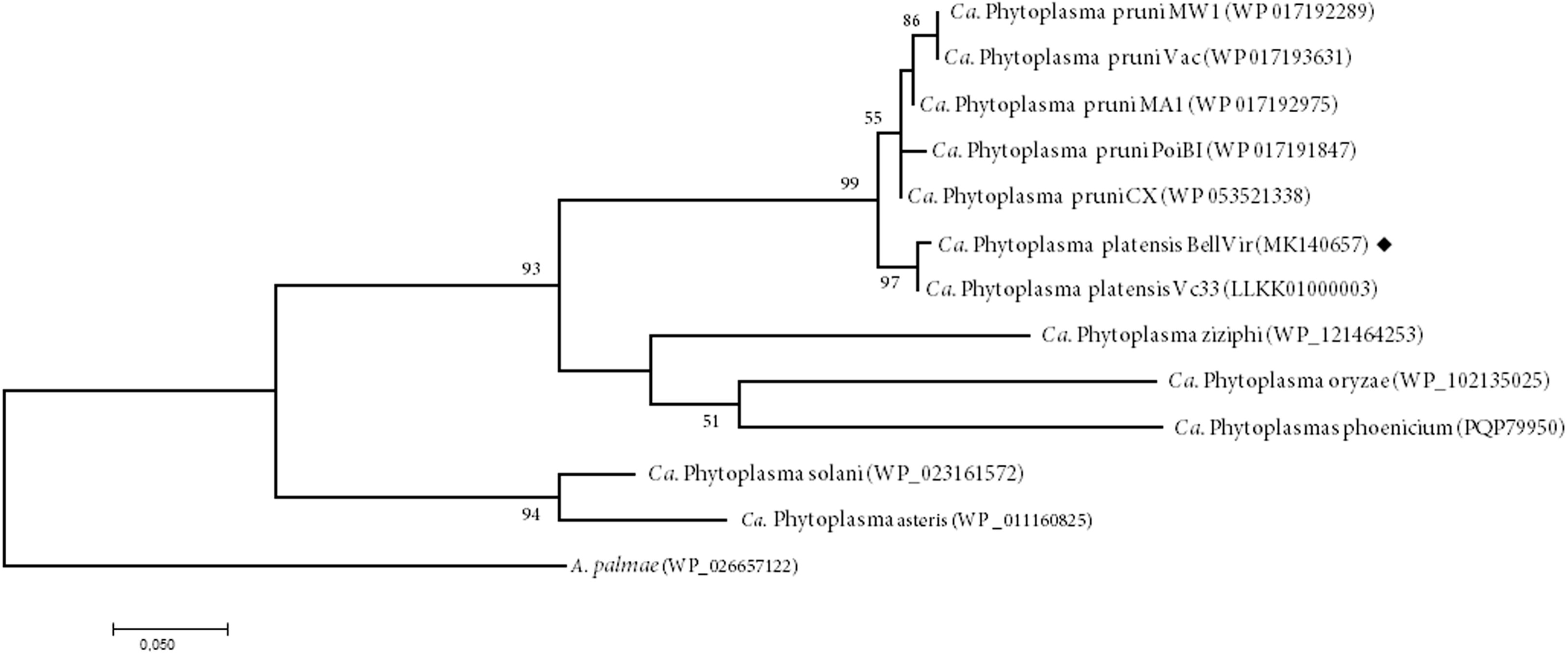
Phylogenetic tree inferred from secA aa sequence analysis using the Maximum Likelihood method implemented in the Molecular Evolutionary Genetics Analysis program (MEGA 6). The numbers on the branches are bootstrap (confidence) values (expressed as percentage of 1000 replicates). GenBank accession number for each taxon is given between parentheses. Sequence of ‘Ca. platensis’ obtained in this work is marked with black diamond. Bar, number of nucleotide substitutions per site.

Based on multilocus sequence analyses, specific vector transmission and geographical distribution, we propose the recognition of the new phytoplasma species ‘*Ca*. Phytoplasma platensis’, within the X-disease clade. The continuous advance in the field of genomics and the fundamental biology of phytoplasmas will allow us to describe new phytoplasma species.

### Description of ‘*Candidatus* Phytoplasma platensis’

‘*Candidatus* Phytoplasma platensis’ (pla. ten’. sis. L. masc. adj. referring to Río de la Plata, a river representative of Argentina and southern South America, where the reference strain was identified).

[(Mollicutes) NC; NA; O, wall less; NAS (GenBank accession number XXXX), oligonucleotide sequences of unique regions of the 16S rRNA gene; 5’-617-CTATAGAAACTGTTTTACTAGAGTGAGTTAGAGGCAAG-654-3’ (*Bellis perennis*, phloem); M].

### Funding information

This work was supported by INTA (PNPV. PE1. 1135022; PNFru; PE2-1105073; PN CI 1108071-1108072); FONCyT PICT 2014-2220 and PICT 2016-0862.

### Ethical statement

No humans or animals were subjects in this work.

