## Supplementary material for "‘Candidatus Phytoplasma platensis’, a novel taxon associated with daisy (Bellis perennis) virescence and related diseases in South America": Table 1

| ***Ca*. Phytoplasma**  **Table 1:** Analysis of 16S rRNA nucleotide identity between ‘Candidatus Phytoplasma platensis’ (BellVir reference strain, MK135798) and ‘Ca. Phytoplasma’ species accepted or proposed to date. | **16Sr*** | **#Genbank** | **Id%** |
| --- | --- | --- | --- |
| *Ca*. Phytoplasma asteris' | 16SrI | M30790 | 90,07 |
| *Ca*. Phytoplasma aurantifolia' | 16SrII | U15442 | 91,13 |
| *Ca*. Phytoplasma australasiae' | 16SrII | Y10097 | 91,44 |
| ***Ca*. Phytoplasma pruni PX11Ct1' rrnA** | **16SrIII** | **JQ044393** | **98,87** |
| ***Ca*. Phytoplasma pruni PX11Ct1' rrnB** | **16SrIII** | **JQ044392** | **98,79** |
| *Ca*. Phytoplasma phoenicium' | 16SrIX | AF515636 | 93,24 |
| *Ca*. Phytoplasma balanitae' | 16SrV | AB689678 | 92,99 |
| *Ca*. Phytoplasma ziziphi' | 16SrV | AB052876 | 93,22 |
| *Ca*. Phytoplasma rubi' | 16SrV | AY197648 | 93,08 |
| *Ca*. Phytoplasma ulmi' | 16SrV | AY197655 | 93,24 |
| *Ca*. Phytoplasma sudamericanum' | 16SrVI | GU292081 | 93,00 |
| *Ca*. Phytoplasma trifolii' | 16SrVI | AY390261 | 93,57 |
| *Ca*. Phytoplasma americanum' | 16SrVI | DQ174122 | 90,05 |
| *Ca*. Phytoplasma fraxini ' | 16SrVII | AF092209 | 92,60 |
| *Ca*. Phytoplasma luffae rrnB' | 16SrVIII | AF353090 | 92,84 |
| *Ca*. Phytoplasma luffae rrnA' | 16SrVIII | AF248956 | 92,84 |
| *Ca*. Phytoplasma spartii' | 16SrX | X92869 | 90,29 |
| *Ca*. Phytoplasma prunorum' | 16SrX | AJ542544 | 90,61 |
| *Ca*. Phytoplasma mali' | 16SrX | AJ542541 | 90,69 |
| *Ca*. Phytoplasma pyri' | 16SrX | AJ542543 | 90,78 |
| *Ca*. Phytoplasma cirsii' | 16SrXI | KR869146 | 94,20 |
| *Ca*. Phytoplasma oryzae' | 16SrXI | AB052873 | 93,62 |
| *Ca*. Phytoplasma convolvuli' | 16SrXII | JN833705 | 89,80 |
| Ca. Phytoplasma fragariae' | 16SrXII | DQ086423 | 90,20 |
| *Ca*. Phytoplasma japonicum' | 16SrXII | AB010425 | 90,12 |
| *Ca*. Phytoplasma australiense' | 16SrXII | L76865 | 89,87 |
| *Ca*. Phytoplasma solani' | 16SrXII | AF248959 | 89,39 |
| *Ca*. Phytoplasma hispanicum' | 16SrXIII | AF248960 | 90,20 |
| *Ca*. Phytoplasma meliae' | 16SrXIII | KU850940 | 90,21 |
| *Ca*. Phytoplasma cynodontis' | 16SrXIV | AJ550984 | 93,96 |
| *Ca*. Phytoplasma castaneae' | 16SrXIX | AB054986 | 92.50 |
| *Ca*. Phytoplasma brasiliense' | 16SrXV | AF147708 | 90,96 |
| *Ca*. Phytoplasma graminis' | 16SrXVI | AY725228 | 86,69 |
| *Ca*. Phytoplasma caricae' | 16SrXVII | AY725234 | 85,73 |
| *Ca*. Phytoplasma rhamni' | 16SrXX | X76431 | 90,71 |
| *Ca*. Phytoplasma pini' | 16SrXXI | AJ632155 | 93,16 |
| *Ca*. Phytoplasma palmicola' | 16SrXXII | EU549768 | 92,44 |
| *Ca*. Phytoplasma omanense' | 16SrXXIX | EF666051 | 92,93 |
| *Ca*. Phytoplasma tamaricis' | 16SrXXX | FJ432664 | 92,93 |
| *Ca*. Phytoplasma costaricanum' | 16SrXXXI | HQ225630 | 90,20 |
| *Ca*. Phytoplasma malaysianum' | 16SrXXXII | EU371934 | 93,24 |
| *Ca*. Phytoplasma wodyetiae' | 16SrXXXVI | KC844879 | 91,85 |
| *Ca*. Phytoplasma noviguineense' | 16SrXXXVII | LC228755 | 92,77 |
| *Ca*. Phytoplasma allocasuarinae' | ND | AY135523 | 90,29 |
| *Ca*. Phytoplasma lycopersici' | ND | EF199549 | 87,05 |
| *Incidentally cited* |  |  |  |
| *Ca*. Phytoplasma palmae' | 16SrIV | U18747 | 93,40 |
| *Ca*. Phytoplasma cocostanzaniae' | 16SrIV | X80117 | 93,96 |
| *Ca*.Phytoplasma vitis' | 16SrV | AF176319 | 93,15 |
