## Supplementary material for "‘Candidatus Phytoplasma platensis’, a novel taxon associated with daisy (Bellis perennis) virescence and related diseases in South America": Table S2

| **Isolate** | **16Sr** | **Genbank** | 283 | 403 | 714 | 732 | 786 | 807 | 810 | 828 | 855 | 897 | 957 | 963 | 981 | 1134 | 1152 | 1162 | 1182 | Country |
| --- | --- | --- | --- | --- | --- | --- | --- | --- | --- | --- | --- | --- | --- | --- | --- | --- | --- | --- | --- | --- |
| **BellVir** | **III-J** | **KC412019** | **A** | **A** | **A** | **C** | **G** | **G** | **G** | **C** | **A** | **C** | **C** | **C** | T | **C** | **T** | **G** | G | Argentina |
| **GDIII** | **III-J** | **KC412014** | **A** | **A** | **A** | **C** | **G** | **G** | **G** | **C** | **A** | **C** | **C** | **C** | T | **C** | **T** | **G** | G | Argentina |
| **TomRed** | **III-J** | **KC412023** | **A** | **A** | **A** | **C** | **G** | **G** | **G** | **C** | **A** | **C** | **C** | **C** | T | **C** | **T** | **G** | G | Argentina |
| **CucVir** | **III-J** | **KC412017** | **A** | **A** | **A** | **C** | **G** | **G** | **G** | **C** | **A** | **C** | **C** | **C** | T | **C** | **T** | **G** | G | Argentina |
| **CassDWB-Arg3** | **III-J** | **MF563965** | **A** | **A** | **A** | **C** | **G** | **G** | **G** | **C** | **A** | **C** | **C** | **C** | T | **C** | **T** | **G** | G | Argentina |
| **CassDWB-Arg4** | **III-J** | **MF563966** | **A** | **A** | **A** | **C** | **G** | **G** | **G** | **C** | **A** | **C** | **C** | **C** | T | **C** | **T** | **G** | G | Argentina |
| **LeSt-Br03** | **III-J** | **KT588312** | **A** | **A** | **A** | **C** | **G** | **G** | **G** | **C** | **A** | **C** | **C** | **C** | T | **C** | **T** | **G** | G | Brazil |
| MxXPV-3 | III-J | KY778014 | G | G | G | T | A | A | A | A | G | T | T | T | C | - | - | - | A | Mexico |
| MxXPV-2 | III-J | KY778015 | G | G | G | T | A | A | A | A | G | T | T | T | C | - | - | - | A | Mexico |
| MxXPV-1 | III-J | KY778016 | G | G | G | T | A | A | A | A | G | T | T | T | C | - | - | - | A | Mexico |
| PX11CT1 | III-A | JQ360960 | G | G | G | T | A | A | A | A | G | T | T | T | C | T | C | T | A | USA |
| PX11CT2 | III-A | JQ360959 | G | G | G | T | A | A | A | A | G | T | T | T | C | T | C | T | A | USA |
| PX92CT1 | III-A | JQ360958 | G | G | G | T | A | A | A | A | G | T | T | T | C | T | C | T | A | USA |
| PX92CT4 | III-A | JQ360957 | G | G | G | T | A | A | A | A | G | T | T | T | C | T | C | T | A | USA |
| CX-95 | III-A | JQ360955 | G | G | G | T | A | A | A | A | G | T | T | T | C | T | C | T | A | USA |
| WX95 | III-A | JQ360956 | G | G | G | T | A | A | A | A | G | T | T | T | C | T | C | T | A | USA |
| PBT | III-C | EF186807 | T | G | G | T | A | A | A | A | G | T | T | T | C | T | C | T | A | USA |
| AKpot6 | III-N | JQ044381 | T | G | G | T | A | A | A | A | G | T | T | T | C | T | C | T | A | USA |
| MT117 | III-M | JQ044380 | T | G | G | T | A | A | A | A | G | T | T | T | C | T | C | T | A | USA |
| VacWB | III-F | EF186809 | T | G | G | T | A | A | A | A | G | T | T | T | C | T | C | T | A | Germany |
| GR1 | III-D | EF186810 | T | G | G | T | A | A | A | A | G | T | T | T | C | T | C | T | A | USA |
| MW1 | III-F | EF186808 | T | G | G | T | A | A | A | A | G | T | T | T | C | - | - | - | A | USA |
| CYE-C | III-B | JQ360962 | G | G | G | T | A | A | A | A | G | T | T | T | C | T | C | T | A | Canada |
| SP1 | III-E | EF186806 | G | G | G | T | A | A | A | A | G | T | T | T | C | T | C | T | A | USA |
| PoiBI | III-H | EF186811 | G | G | G | T | A | A | A | A | G | T | T | T | C | T | C | T | A | USA |
| WWB | III-G | JQ360961 | G | G | G | T | A | A | A | A | G | T | T | T | C | T | C | T | A | USA |

**Table S2**: Single Nucleotide Polimorphism (SNP) in ribosomal protein sequence that distinguish ‘*Ca*. Phytoplasma platensis’ (bold) strains from ‘*Ca*. phytoplasma pruni’ strains
