## Supplementary material for "‘Candidatus Phytoplasma platensis’, a novel taxon associated with daisy (Bellis perennis) virescence and related diseases in South America": Table S3

**Table S3:** List of primers used in this work. *: expected amplified fragment size

| **Primer** | **Sequence** | **Ann/Tº** | **bp*** | **Reference** |
| --- | --- | --- | --- | --- |
| P1 | AAGAGTTTGATCCTGGCTCAGGATT | 55ºC | 1800 | Deng and Hiruki, 1991 |
| P7 | CGTCCTTCATCGGCTCTT |  |  |  |
| secAfor1 | GARATGAAAACTGGRGAAGG | 60ºC | 840 | Hodgetts et al., 2008 |
| secARv | GTTTTRGCAGTTCCTGTCATNCC |  |  |  |
| rpF1 | GGACATAAGTTAGGTGAATTT | 54ºC | 1300 | Lim and Sears, 1992 |
| rpR1 | ACGATATTTAGTTCTTTTTGG |  |  |  |
| Imp-XdFw1 | ATCTCGTCCTCTTAAACCGCATCC | 54ºC | 1000 | This paper |
| Imp-XdRv1 | AGACTCTTAACTGGCAACG |  |  |  |
| idpA-XdFw1 | CCCTTCTGCTCCGCCAATTA | 56ºC | 1400 | This paper |
| idpA-XdFw1 | ATTGTTCTTTTTCTCGGCAA |  |  |  |
