## Supplementary material for "‘Candidatus Phytoplasma platensis’, a novel taxon associated with daisy (Bellis perennis) virescence and related diseases in South America"

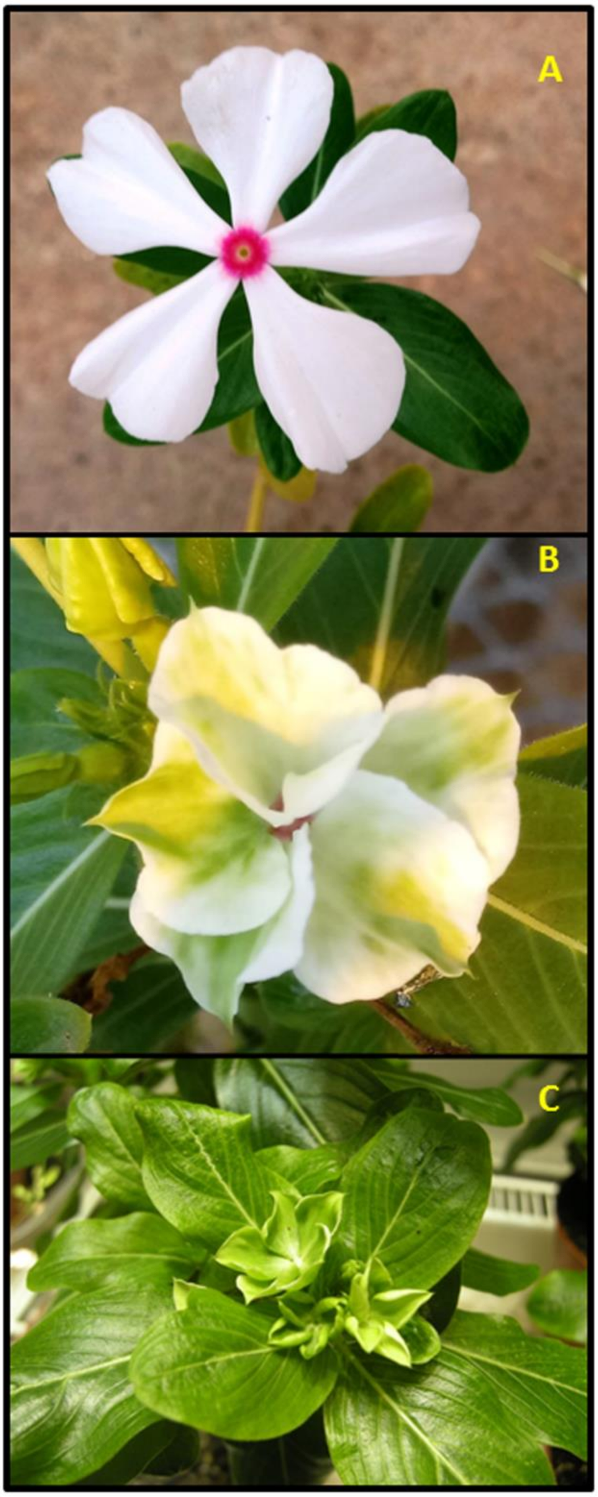


**Figure S1:** BellVir phytoplasma infection caused symptoms of virescence (B) and phyllody (C) on periwinkle grafted plants. A: asymptomatic plant

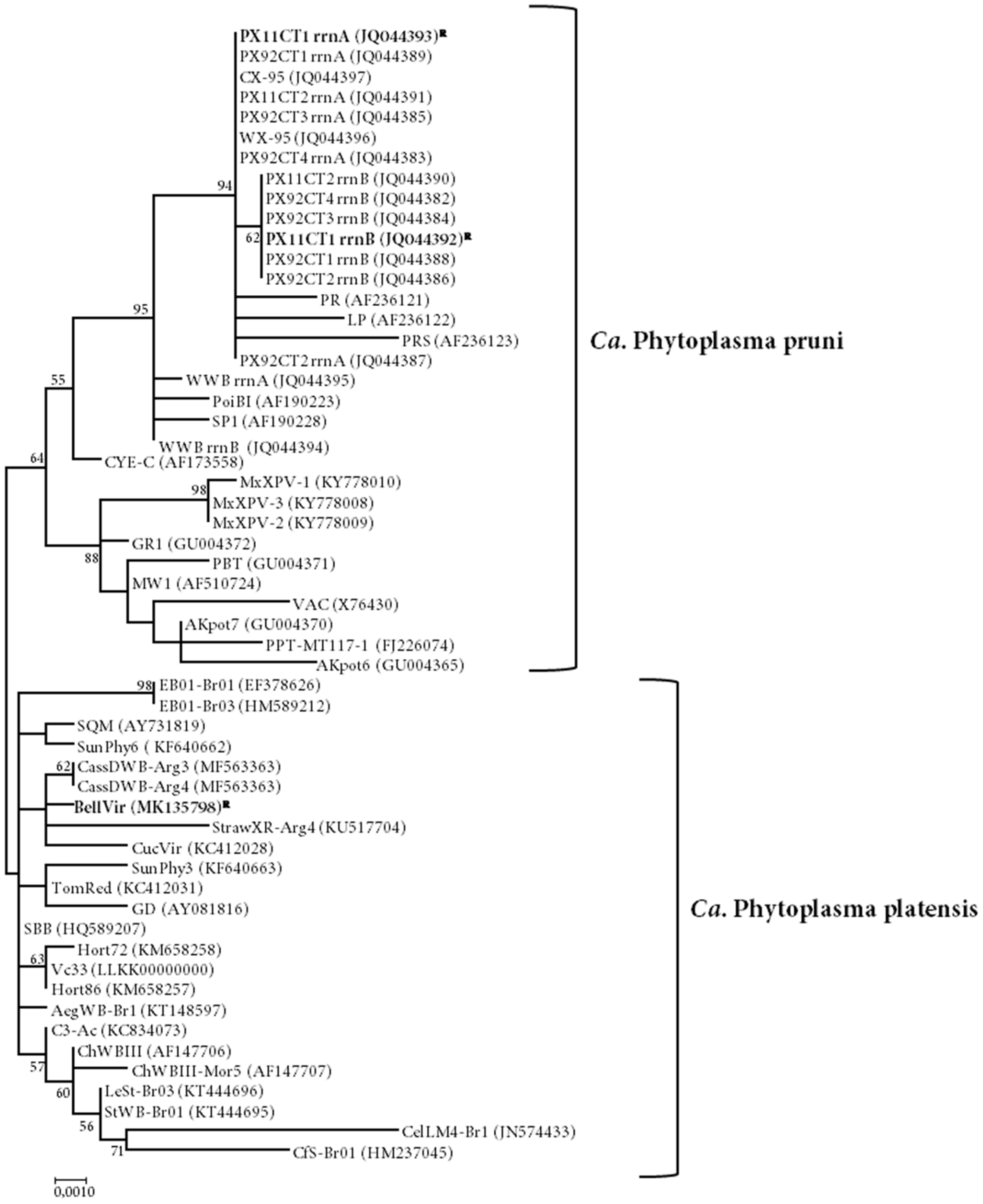


**Figure S2:** Phylogenetic tree inferred from analysis of 16S rDNA sequences using the Maximum-Likelihood method. The numbers on the branches are bootstrap (confidence) values (expressed as percentage of 1000 replicates). R: reference strains

**#1**
pruni 5’-CACATTAGTTAGTTGG**T**AGGGTAAAGGCCTACC-3’
platensis 5’-CACATTAGTTAGTTGG**C**AGGGTAAAGGCCTACC-3’

**#2**
pruni 5’-TAATAAGTCTATAGTTTAATTTCAG**T**GCTTAACGCTGTTGTGCTATAG-3’
platensis 5’-TAATAAGTCTATAGTTTAATTTCAG**C**GCTTAACGCTGTTGTGCTATAG-3’

**#3**
pruni 5’- TAAAAC**T**GGTAC -3’
platensis 5’-TAAAAC**C**GGTAC -3’

**#4**
pruni 5’-ATGGAGGT**C**ATCAGGAAAACAGGTGGTGC-3’
platensis 5’-ATGGAGGT**T**ATCAGGAAAACAGGTGGTGC-3’

**#5**
pruni 5’-CTTGTCGTTA**G**TTGCCAGCATGTAAT-3’
platensis 5’-CTTGTCGTTA**A**TTGCCAGCATGTAAT-3’

**#6**
pruni 5’-GATGGGGACTTTAA**C**GA-3’
platensis 5’-GATGGGGACTTTAA**T**GA-3

**#7**
pruni 5’-TCTCA**A**AAAATCAATC-3’
platensis 5’-TCTCA**C**AAAATCAATC-3’

**Figure S3:** Aligned nucleotide sequence of 16S rRNA showing the 7 out 13 unique features of ‘*Ca*. phytoplasma pruni’ vs same region of ‘*Ca*. Phytoplasma platensis’. The asterisk shows the presence of nucleotide differences

a

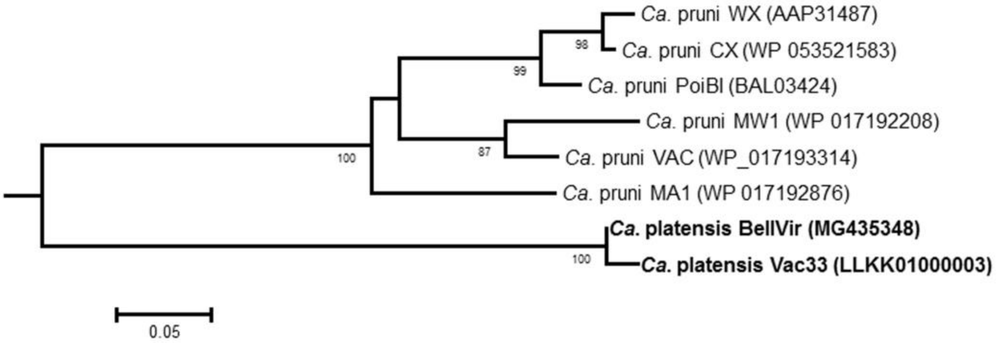


b

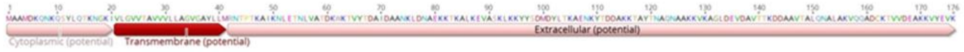


c

| | BellVir | Vc33 | PoiBI | WX | CX | MA1 | MW1 | VAC |
|---|---|---|---|---|---|---|---|---|
| BellVir | | 97.73% | 58.33% | 58.43% | 58.99% | 60.11% | 59.32% | 61.02% |
| Vc33 | 97.73% | | 56.67% | 57.87% | 58.43% | 59.55% | 58.76% | 60.45% |
| PoiBI | 58.33% | 56.67% | | 92.70% | 93.26% | 82.02% | 79.66% | 82.49% |
| WX | 58.43% | 57.87% | 92.70% | | 97.73% | 82.39% | 81.14% | 82.29% |
| CX | 58.99% | 58.43% | 93.26% | 97.73% | | 81.25% | 81.14% | 82.86% |
| MA1 | 60.11% | 59.55% | 82.02% | 82.39% | 81.25% | | 80.00% | 83.43% |
| MW1 | 59.32% | 58.76% | 79.66% | 81.14% | 81.14% | 80.00% | | 90.29% |
| VAC | 61.02% | 60.45% | 82.49% | 82.29% | 82.86% | 83.43% | 90.29% | |

**Figure S4:** Analysis of imp aa sequence reveals difference between '*Ca*. pruni' and '*Ca*. platensis'. a: Phylogenetic tree based on imp aa sequence of *Ca*. pruni' and '*Ca*. platensis' (bold) strains. The numbers on the branches are bootstrap (confidence) values (expressed as percentage of 1000 replicates).b: Structure of imp protein from 'Ca. platensis' showing typical transmembrane domain at the N-terminal and the extracellular at C-terminal. c: aa %identity values

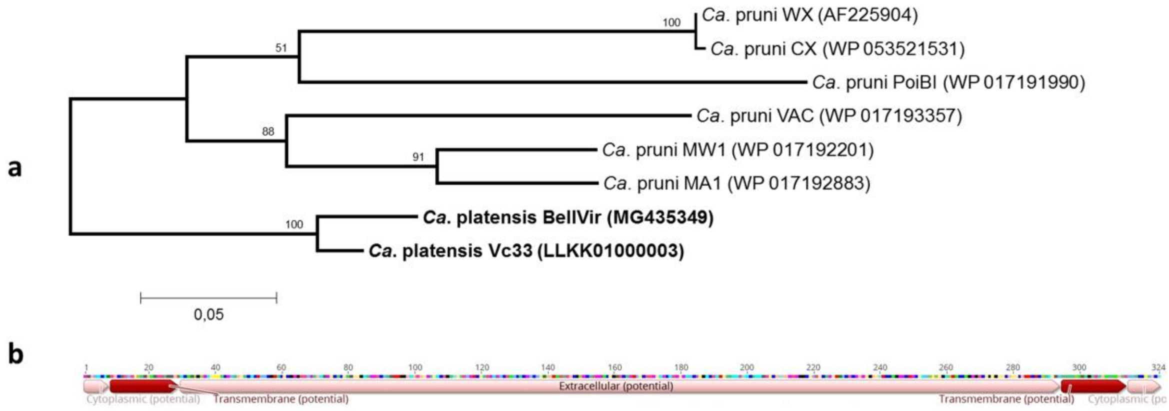


c

| | BellVir | Vc33 | PoiBI | VAC | MW1 | MA1 | WX | CX |
|---|---|---|---|---|---|---|---|---|
| BellVir | | 87.04% | 72.26% | 70.25% | 69.51% | 68.31% | 66.26% | 65.95% |
| Vc33 | 87.04% | | 68.90% | 69.33% | 69.21% | 75.58% | 70.72% | 70.39% |
| PoiBI | 72.26% | 68.90% | | 71.04% | 71.60% | 68.90% | 64.74% | 64.44% |
| VAC | 70.25% | 69.33% | 71.04% | | 76.60% | 72.70% | 63.61% | 63.61% |
| MW1 | 69.51% | 69.21% | 71.60% | 76.60% | | 81.71% | 66.26% | 66.26% |
| MA1 | 68.31% | 75.58% | 68.90% | 72.70% | 81.71% | | 73.03% | 73.03% |
| WX | 66.26% | 70.72% | 64.74% | 63.61% | 66.26% | 73.03% | | 99.65% |
| CX | 65.95% | 70.39% | 64.44% | 63.61% | 66.26% | 73.03% | 99.65% | |

**Figure S5:** Analysis of idpA aa sequence reveals difference between '*Ca*. pruni' and '*Ca*. platensis'. a: Phylogenetic tree based on idpA aa sequence of *Ca*. pruni' and '*Ca*. platensis' (bold) strains. The numbers on the branches are bootstrap (confidence) values (expressed as percentage of 1000 replicates).b: Structure of imp protein from 'Ca. platensis' showing typical transmembrane domain at the N-terminal and C—Terminal while the extracellular domain remains at the center. c: aa %identity values
